# Gigantic Macroraptorial Sperm Whale Tooth (cf. *Livyatan*) from the Miocene of Orange County, California

**DOI:** 10.1101/2023.01.25.525567

**Authors:** Kristin I. Watmore, Donald R. Prothero

**Affiliations:** Department of Geological Sciences, California State Polytechnic University, Pomona, CA 91768; Department of Vertebrate Paleontology, Natural History Museum of Los Angeles County, Los Angeles, California USA 90041

## Abstract

A fossil tooth from the middle-upper Miocene Monterey or Capistrano Formation in the Orange County Cooper Center Paleontology Collection shows that a gargantuan sperm whale once inhabited the Miocene seas of southern California. Though difficult to diagnose to genus level based on a single, incomplete tooth, comparisons with known Miocene physeteroid whales provide key insight into the affinities of this fossil. Even though the tip is broken, the entire tooth is over 250 mm long and 86 mm in diameter. It has enamel only on the tip of the broken crown and there is no enamel coating over the rest of the tooth. Instead, the tooth consists mostly of layers of cementum over a core of ossified dentin. It is slightly smaller than the largest teeth of the largest known physeteroid, the South American Miocene *Livyatan melvillei*, a genus that has not been found outside of the Southern Hemisphere besides one record from northern Europe. It is also just slightly smaller than similar gigantic teeth reported from South Africa and Australia from the middle-late Miocene through the early Pliocene. We compared it to other Miocene members of the family such as *Hoplocetus*, *Scaldicetus*, and *Zygophyseter*, but none have teeth as large as this one. It is bigger than all known specimens of all other North American Miocene physeteroid whales, including another whale from the Monterey Formation, *Albicetus oxymycterus*. This fossil suggests that giant physeteroid whales closely related to *Livyatan* lived in the North Pacific. It represents a substantial geographic range extension for giant physeteroid whales, previously known only from the Miocene of the Southern Hemisphere and northern Europe.

## Introduction

The sperm whales (family Physeteridae) are represented today by three species, *Physeter macrocephalus*, the giant sperm whale and two species of pygmy sperm whale, *Kogia breviceps* and *K. sima*. They are found throughout the world’s oceans today, but have an even more extensive fossil record, with at least 20 genera going back to the late Oligocene (recently reviewed by [1]). Additional questionable genera are represented by isolated fragmentary specimens, including teeth.

According to Velez-Juarbe et al. [2], the modern Physeteridae are a relatively derived branch of the stem-group they called the Pan-Physeteroidea. More basal members of the clade include a variety of sperm whales with very different features and sizes. The most spectacular was the giant macroraptorial whale *Livyatan melvillei* [3,4], which may have reached 13-17 meters in total body length. It had large piercing teeth in both the upper and lower jaws up to 362 mm long, making them the largest non-tusk teeth known in any mammal. The teeth equipped the animal for catching large prey like other whales and sharks, as well as smaller animals. In contrast, the modern sperm whale, *Physeter macrocephalus*, has no upper teeth and uses deep-sea diving to prey on giant squid. As discussed below, teeth possibly referable to *Livyatan* have been reported from Peru, Argentina, and South Africa and Australia, as well as the North Sea. This is the first specimen reported from North America and the North Pacific realm.

## Materials and Methods

Measurements were made with digital dial calipers and for the longer dimensions or circumferences, a flexible metric measuring tape was used.

The specimen, OCPC 3125/66099 (Figs 1,2) comes from the Mission Viejo housing development in southern Orange County, California. Unfortunately, the people who were doing salvage paleontology at the site in the 1980s and 1990s kept very poor field records, and its precise geographic location and stratigraphic level in the development cannot be determined. It had a field number MV60 painted on it, which suggests that it comes from planning area 28, which has both Monterey and Capistrano formations (middle to upper Miocene) exposed in the area. Although some specimen labels suggest it might have come from the middle-upper Miocene Monterey Formation, the preservation of the tooth more closely resembles that of the Capistrano Formation fossils. The Capistrano Formation is uppermost Miocene (about 6.6-5.8 Ma), according to Barboza et al. [5]. Barboza et al. [5] mentioned the occurrence of two different unidentified species of physeteroids from the Oso Member of the Capistrano Formation, but no further details were ever provided. It is not clear what these identifications were based on or whether OCPC 3125/66099 was included among the two physeteroids reported in that paper.

**Fig 1.**
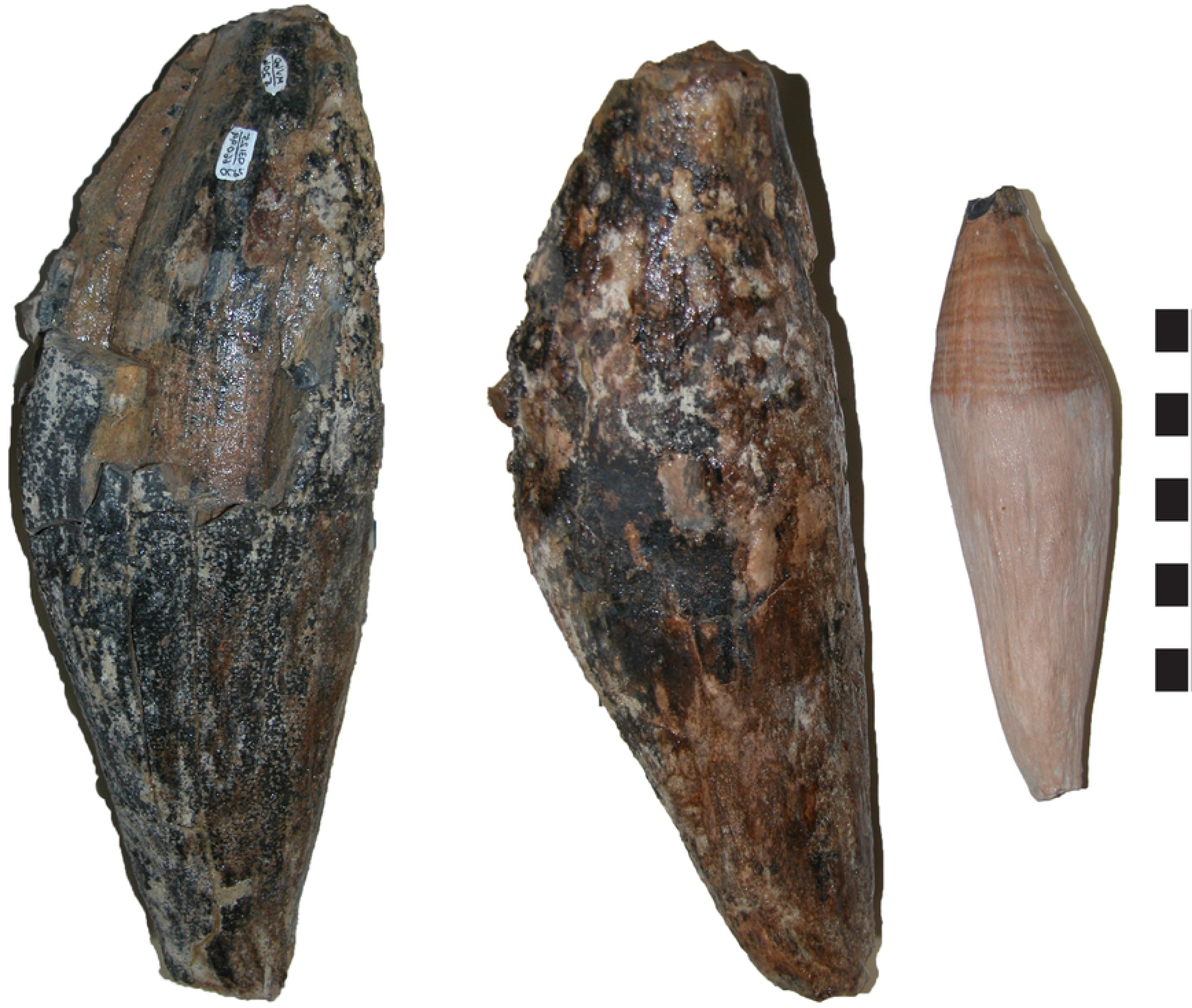
Two views of OCPC 3125/66099 (left) compared with a modern sperm whale tooth (*Physeter macrocephalus*) tooth on the right. Scale bar in cm.

**Fig 2.**
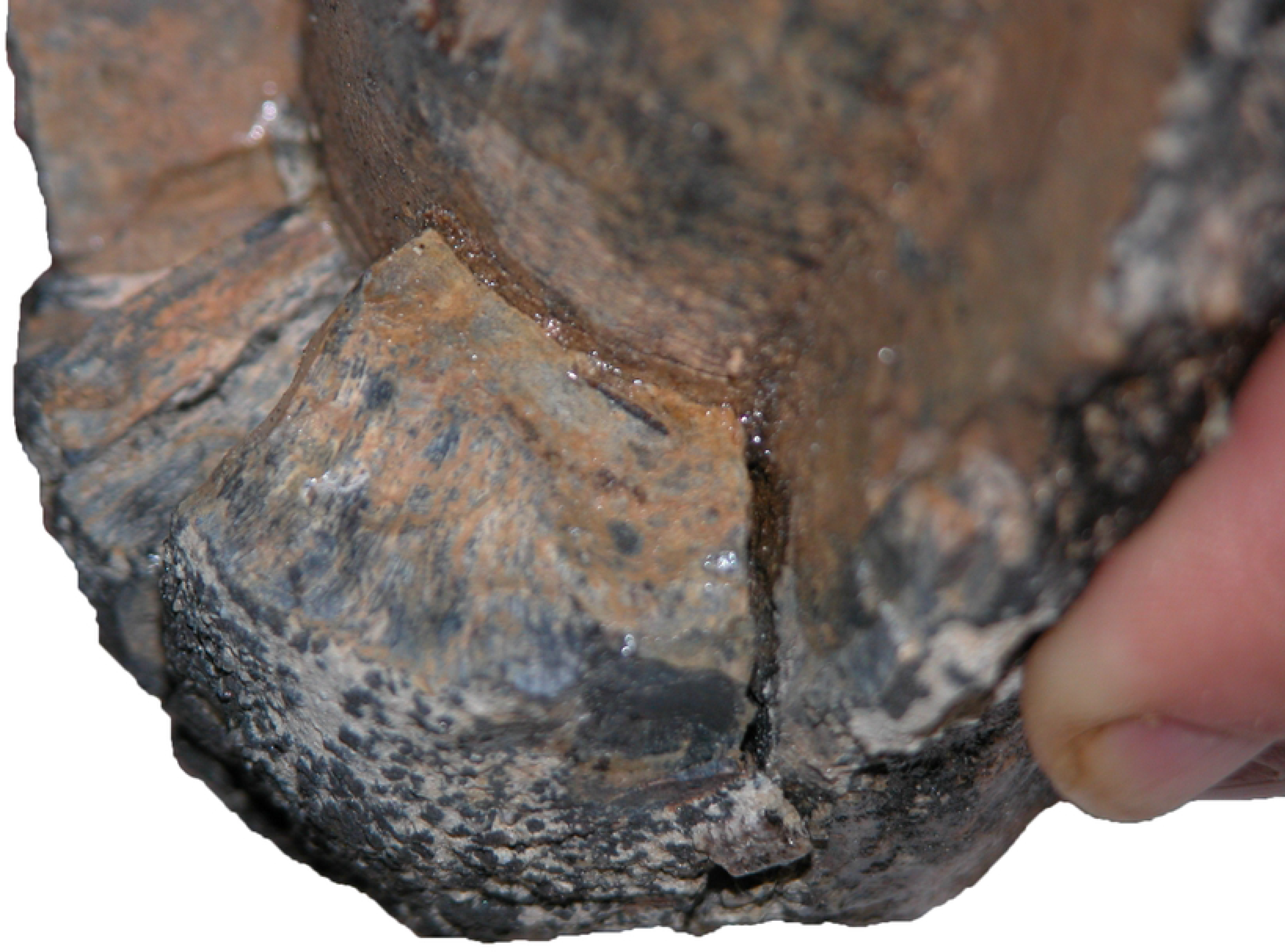
Close up of the broken crown of the tooth, showing the thick layer of outer cementum and the core of ossified dentin.

## Results

OCPC 3125/66099 (Fig 1) consists of a single isolated tooth with a single root and crown, which taper dramatically from the middle of the tooth in both crown and root directions, a condition typical of physeteroid whales. Its minimum total length is currently 250 mm, although the broken tips of both the crown and the root of the tooth suggest it was a bit longer. Its diameter is 86 mm at the widest point, and the cross-section in the middle of the tooth is circular, with a circumference of 252 mm at its widest point. The base of the broken crown of the tooth is oval in cross-section, measuring 29 mm by 36.5 mm in dimensions. There was apparently a tiny tip of enamel on the crown of the tooth, although it is broken off and only a few remnants of enamel are found at the broken edge. At its thickest spot, the enamel is only 3-4 mm thick. Unlike the teeth of *Livyatan* [3,4], there was apparently no enamel coating over the rest of the crown. Because OCPC 3125/66099 did not have a coating of enamel over most of the tooth, it is not possible to determine if there were any marks of wear on the surface. In addition, the surface of the tooth is weathered, so any surface marks would have been destroyed anyway. The bulk of the tooth is composed of a thick layer of cementum, over a core of ossified dentin. On one side of the crown of the tooth, the outer cementum layer is broken away (Fig 2), exposing the core of ossified dentin. The thickness of the outer cementum layer measures 25 mm, so the core of ossified dentin was about 36 mm in diameter in this portion of the tooth.

## Discussion

Though this fossil is incomplete, there are valuable comparisons that can be made between it and other known fossil physeteroid whales. The most similar known fossils come from the enormous macroraptorial sperm whale from southern Peru described by Lambert et al. [3], named *Livyatan melvillei* [3,4]. Our tooth conservatively measures approximately 100 mm shorter in total length compared to the largest *Livyatan melvillei* tooth (Fig 3). However, the shortest tooth of *Livyatan melvillei* reported by Lambert et al. [3] is approximately 115 mm in total length, which is about 135 mm shorter than our tooth, and our tooth lacks a crown and part of the root which could add several centimeters to the length [3]. In diameter, our tooth falls within the lower part of the range of *Livyatan* diameters, which span between 81–121 mm [3]. It is worth noting that apices of the lower teeth in *Livyatan melvillei* are more recurved than those of the posterior teeth, which exhibit a more cylindrical shape. Similarly, our tooth displays that more circular cross-section shape around the tooth midsection. Interestingly, our tooth is 5 mm longer than the apical tooth of *Livyatan melvillei* and less recurved, suggesting our tooth is more posteriorly positioned. In terms of cementum thickness, our tooth falls within the range of *Livyatan melvillei* cementum thickness of 21–28 mm thick [3]. In Argentina, similar physeteroid teeth were described [6] with maximum lengths and diameters less than those respective measurements in both *Livyatan melvillei* from Peru, and the California specimen (Fig 4), yet the Argentinian material was still classified as belonging to the genus *Livyatan*. Specifically, the teeth from Argentina measured in length between 142–178 mm (smaller than the California tooth at 250 mm), and diameter between 72-74 mm. This puts the Argentinian *Livyatan* at 72 mm shorter than our tooth in length and 12 mm narrower in width (Fig 4). Morphologically, our specimen shares distinctive features with the *Livyatan* teeth described by both Lambert et al. [3,4] and Piazza et al. [6], including the robust mid-section of the tooth adjoined to the gradual tapering and bending towards both the root and crown ends. *Livyatan melvillei* was also reported from the upper Miocene Bahía Inglesa Formation in Chile [7], although the measurements of the teeth were not published.

**Fig 3.**
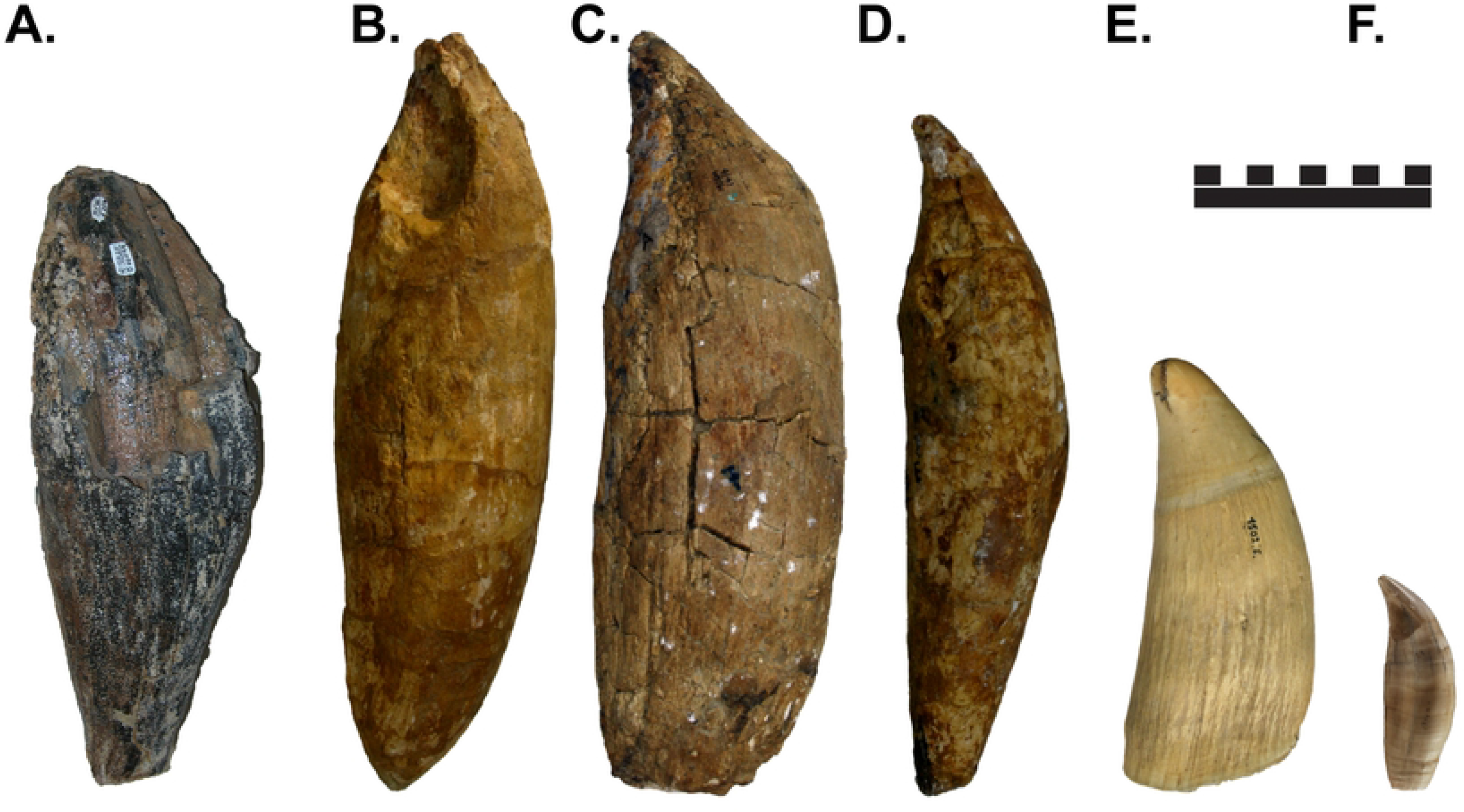
Comparison of (A) OCPC 3125/66099 with several teeth of *Livyatan melvillei* from Peru (B-D), as well as a modern sperm whale (*Physeter macrocephalus*) tooth (E), and a killer whale (*Orcinus orca*) tooth (F). Largest *Livyatan* teeth measure about 360 mm in total length. Scale bar in cm. (B-E courtesy O. Lambert).

**Fig 4.**
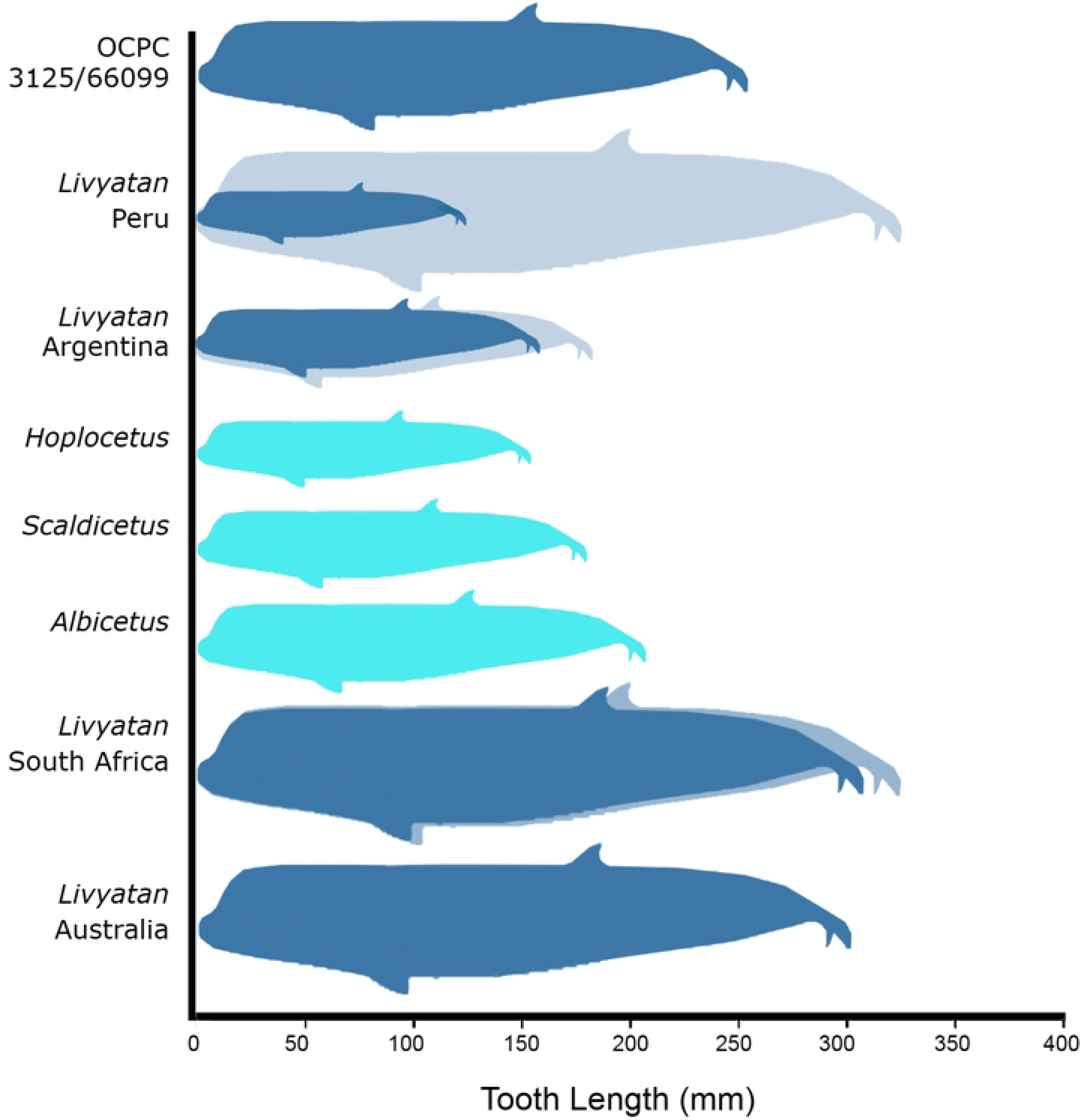
Visual comparison of the range of sizes of OCPC 3125/66099 (top) with the minimum and maximum lengths of other known teeth of *Livyatan*, and of other extinct physeteroid whales. Minimum and maximum sizes shown by the different sized silhouettes. Non-*Livyatan* whales are shown in light blue (Original art by K. Marriott).

Besides the fossils from South America, most other specimens of the size of *Livyatan* are reported from the Southern Hemisphere (Fig 4). Two *Livyatan* teeth were reported from the late Miocene of South Africa [8] from Pliocene deposits near the Hondeklip Bay village of Namaqualand. These specimens are stored in the Iziko South African Museum. They are cataloged as SAM-PQHB-433 and SAM-PQHB-1519, and measure 325 millimeters and 301 millimeters in height respectively, the latter specimen missing its crown. The larger tooth is longer than the California tooth by about 75 mm. In addition, another giant physteroid tooth measuring 300 mm in length (about 50 mm longer than the California tooth) were reported from Pliocene deposits in Beaumaris Bay, near Melbourne, Australia, but it has never been formally described [9]. It is now in the National Museums Victoria, and bears the catalogue number NMV P16205.

Another genus of notable comparison to our specimen is *Albicetus*. Collected from the Monterey Formation in Santa Barbara California, *A. oxymycterus* teeth measure approximately 205 mm in length, 80 mm in diameter, have a thick cementum layer, midsection robustness with tapering towards tips and a posterior curvature moving distally up the tooth [1]. The length measurement is around 45 mm less than our tooth and the width measurement is approximately 6 mm smaller. It should be noted that, phylogenetically, Boersma and Pyenson [1] placed *Albicetus* in an unresolved polytomy with *Livyatan* and described it as a smaller, but morphologically similar whale.

Other genera such as *Scaldicetus*, *Zygophyseter, Brygmomphyseter, Acrophyseter* and *Hoplocetus* can provide other valuable comparisons (Fig 4). *Scaldicetus* teeth discovered in the Guadalquivir Basin of southern Spain, have conservative total length measurements not exceeding 180 mm and diameter measurements of 18 mm and 19 mm for each respective tooth [10]. The genus *Zygophyseter* is known from a fairly complete specimen from the Cisterna Quarry of southern Italy, with the most anterior teeth measuring longer than the posterior teeth [11]. *Zygophyseter* has larger teeth than *Scaldicetus*, however, the length measurements do not approach the length of our tooth. In terms of overall shape, the anterior teeth of *Zygophyseter* are strongly anterolaterally bent, a condition not seen to the same extent in our tooth (though we cannot determine with confidence the original position of the tooth in the whale’s mouth). *Hoplocetus ritzi* [12] from the Schleswig-Holstein of northern Germany, includes a probable lateral tooth reaching a maximum length of 150 mm and another lateral tooth exceeding 154 mm in circumference, which is much less than the respective measurements of our tooth. Morphologically, the *Hoplocetus* teeth resemble our tooth with the robust midsection, slight curvature, and tapering toward the crown and roots. However, the *Hoplocetus* teeth are smaller than our tooth and are less than 100 mm of the length and 98 mm of circumference of our tooth.

Overall measurements of our specimen fall in between the limits of measurements observed in both Lambert et. al [3,4] and Piazza et. al [6] specimens in Peru, Argentina, and Chile in South America, which were all classified as *Livyatan* (Figs 3,4). Also, the small amount of apparent enamel left on our tooth (a character not seen in modern physeteroid whales) suggests that our tooth represents a basal physeteroid group. Altogether, it seems that our tooth comes from a physeteroid whale of substantial size within range of medium-sized teeth assigned to *Livyatan*. If this is so, it implies a marked range extension for *Livyatan*, as prior descriptions of these animals have been restricted to the Southern Hemisphere, except for a single broken tooth from the North Sea [13]. Previously, several authors [3,4,6] speculated as to why no *Livyatan* had been found in the Northern Hemisphere, with suggestions that it was unable to cross equatorial oceans, or that its absence was possibly due to collection bias. We now know that it was simply luck of collecting these relatively rare fossils. The California fossil firmly establishes that a whale like *Livyatan* swam in the Pacific middle-late Miocene seas of southern California. Thus, *Livyatan* had a virtually global distribution, like many open-marine whales which live in most of the world’s oceans.

*Livyatan* was the apex predator in several middle-late Miocene and Pliocene marine communities around the world. For example, in addition to *Livyatan*, the Pisco Formation yields a wide diversity of different types of marine vertebrates, including baleen whales, additional toothed whales, pinnipeds, marine sloths, penguins and other birds, including the giant soaring sea bird *Pelagornis*, crocodilians, sea turtles, as well as a large diversity of sharks (including *Otodus megalodon*) and other fish. Similarly, the Bahía Inglesa Formation in Chile yields not only *Livyatan*, but also a variety of baleen and toothed whales, pinnipeds, marine sloths, 28 species of shark (including *O. megalodon*), bony fish, penguins and other marine birds, and crocodilians. The Australian *Livyatan* comes from the lower Pliocene Black Rock Sandstone in Beaumaris Bay near Melbourne, Australia, which yields over 20 species of shark (including *O. megalodon*), the physeteroid *Physetodon baileyi* and other whales, elephant seals, dugongs, sea turtles, penguins and a variety of giant soaring sea birds such as *Pelagornis*. The South African teeth come from the Avontuur Member of the Alexander Bay Formation near Hondeklip Bay, Namaqualand, which is close to the upwelling of the Benguela Current, so it also yields many baleen whales, dolphins and other toothed whales, as well as abundant shark teeth (including *O. megalodon*). However, many other famous marine middle-late Miocene and Pliocene localities (such as Sharktooth Hill in California, Lee Creek Mine in North Carolina, and the Chesapeake Bay Miocene) had a similar high diversity of marine mammals, sharks, and fish, but so far *Livyatan* has not been reported from them. Despite its scarcity, however, *Livyatan* was clearly a wide-ranging taxon and performed the apex predator role in many different Miocene and Pliocene ecosystems.

During the early Pliocene, macroraptorial sperm whales vanished alongside many other kinds of whales, other marine mammals, sharks (including *O. megalodon*), and other fish. Most authors (e.g., [3, 14]) attribute this extinction to the rapid global cooling in the Pliocene.

## Conclusions

A tooth referable to the gigantic macropredatory physeteroid whale *Livyatan*, originally known only from South America, South Africa, Australia, and the North Sea, is now reported from the middle-late Miocene of southern California. This establishes that huge *Livyatan*-like physeteroids had a global distribution in the middle Miocene to Pliocene, although they are so rare that they are not found in most marine fossil localities of that age.

## Acknowledgments

We thank W. Gelnaw and M. Madan-Richards for access to the specimen. We thank O. Lambert for providing us the original images of the teeth of *Livyatan*, on which we based Fig 3. We thank E. Prothero for Photoshopping the images, and K. Marriott for drawing Fig 4. We thank O. Lambert and xxxx for reviewing the paper.

## Author Contributions

Both authors contributed equally to all parts of this paper.

